# Diverse Genomic Landscape of Swine Influenza A Virus in England (2014–2021)

**DOI:** 10.1101/2025.04.28.650978

**Authors:** Benjamin C. Mollett, Alexander MP Byrne, Helen E. Everett, Scott M Reid, Susanna Williamson, Tavis K Anderson, Joe James, Ashley C Banyard, Ian H Brown, Nicola S. Lewis

**Affiliations:** Virology, Animal and Plant Health Agency (APHA), New Haw, Addlestone, UK; Pathobiology & Population Sciences, Royal Veterinary College, London, UK; Worldwide Influenza Centre, WHO Collaborating Centre for Reference and Research on Influenza, The Francis Crick Institute, London, UK; The WOAH/FAO International Reference Laboratory for Animal Influenza, Animal and Plant Health Agency (APHA), New Haw, Addlestone, UK; Veterinary Investigation Centres, Animal and Plant Health Agency (APHA), Bury St Edmunds, and Thirsk, UK; Virus and Prion Research Unit, National Animal Disease Centre, Agricultural Research Service, U.S Department of Agriculture Ames, Iowa, IA 50010; The Pirbright Institute, Pirbright, Woking, UK

**Author notes:** Corresponding author by.

## Abstract

Surveillance of influenza A viruses in pigs (SwIAV) is critical for identification of novel genetic groups that pose a risk to pig health and might have zoonotic potential. SwIAVs circulating in pigs in England between 2014 and 2021 were characterised using whole genome sequencing (WGS).

Haemagglutinin (HA) and neuraminidase (NA) sequencing data from 82 of 368 influenza A positive samples (71 submissions) were determined, identifying H1N1 and H1N2 subtypes from the 1A classical swine and 1B human-seasonal lineages respectively. The 1B lineage viruses were predominant, accounting for 68.29% of sequenced viruses, with 1A lineage viruses comprising 31.71%, primarily from the 1A.3.3.2 clade (2009 H1N1 pandemic origin). This study characterised previously undefined diversity within the 1B lineage which led to the designation of new HA clades 1B.1.1.1, 1B.1.1.2 and 1B.1.1.3. Complete genome data were obtained from 64/82 viruses thereby updating the definition of genetic diversity thresholds and leading to the identification of 24 unique genotypes. All these 64 viruses contained PB2, PB1, PA, NP, MP, and NS gene segments of 2009 H1N1 pandemic origin. These data highlight the increasing divergence of SwIAV within pig populations England and emphasise the requirement for continued genomic surveillance to improve animal health and monitor zoonotic risk.

## Introduction

Swine influenza A viruses (SwIAVs) are respiratory pathogens that have a global impact on pig health and welfare and pose a zoonotic risk (Vincent et al., 2020). Infection of pigs with these viruses can result in substantial economic losses, including reduced productivity, expenditure on treatment and control measures, including vaccination, and increased mortality through comorbidity with other diseases (e.g. bacterial and other viral pathogens). SwIAVs also present zoonotic threat with spillovers to humans and were associated with the 2009 H1N1 pandemic (Anderson et al., 2021, Rambo-Martin et al., 2020).

The evolution of SwIAV is complex. As with human-origin influenza A viruses, classification is based on the major antigenic determinants, the haemagglutinin (HA) and neuraminidase (NA) envelope glycoproteins present on the surface of the virion (Lewis et al., 2016, Air, 1981). The lack of RNA polymerase proofreading during the replication cycle of influenza viruses helps to drive continuing diversity, and the segmented genome facilitates reassortment events wherever co-infection occurs (Webster et al., 1982). The genetic diversity of SwIAVs in pigs is strongly influenced by their direct or indirect interfaces with other infected species, especially humans and birds. Human influenza A viruses have been transmitted to pigs on multiple occasions (Nelson and Vincent, 2015). Since the initial detection in 1979 in Europe, the emergence and circulation of avian-origin H1N1 viruses has resulted in a lineage of SwIAVs termed ‘avian-like’ H1N1 (Krumbholz et al., 2014), reclassified as the H1 1C lineage under the global classification system (Anderson et al., 2016). In 1984, the re-emergence of H3N2 viruses in swine occurred, likely through reassortment between a human seasonal H3N2 virus and the ‘avian-like’ H1 1C swine H1N1 virus already circulating in pigs (Simon et al., 2014). In 1994, H1N2 viruses were identified. Their HA gene originated from a human seasonal H1N1 virus that was dominant in the mid-1980s, while the remaining gene segments were acquired from a human-like swine H3N2 virus (Brown et al., 1998). Commonly referred to as ‘human-like’ H1N2 viruses, their HA classification aligns with global H1 lineage 1B (Anderson et al., 2016). As such, prior to the emergence of the H1N1 pandemic virus in 2009, three main subtypes of SwIAV (H1N1, H1N2 and H3N2) were likely circulating in Europe (Zell et al., 2013), but with regional variations, including in Great Britain (GB); H3N2 has not been detected in GB pig populations since 1997 (Brown, 2013). Then, in 2009, a swine origin pandemic (pdm) H1N1 virus emerged globally in human populations involving a virus containing an HA originating from the classical swine H1N1 lineage, globally designated as lineage 1A.3.3.2 (Harder et al., 2013, Watson et al., 2015). This virus has been reintroduced into pigs repeatedly (Nelson and Vincent, 2015, Watson et al., 2015, Markin et al., 2023a) following reverse zoonotic events.

The increased detection of SwIAV infections within the European pig population, coupled with the co-circulation of multiple lineages, has contributed to the emergence of reassortant strains possessing a diverse combination of swine, avian and human genetic segments (Henritzi et al., 2020). In GB, surveillance is predominantly focussed on England which has the highest pig density. Within England, three SwIAV lineages circulate with distinct HA genes: the 1A.3.3.2 (H1pdm), 1B (H1hu) and 1C (H1av) (Anderson et al., 2016). Therefore, the potential for increased genetic diversity exists and, whilst surveillance for these viruses is incomplete, existing data suggests that the antigenic landscape continues to change in England and globally. An example of this is the emergence of pandemic H1 1A.3.3.2 viruses that are seemingly displacing the previously dominant H1 1C viruses (Watson et al., 2015). The viral genetic diversity is further increased by reassortment events involving the other six viral gene segments encoding the polymerase complex; polymerase basic 1 and 2 (PB1 and PB2, respectively), polymerase acidic (PA) and nucleoprotein (NP) segments; matrix proteins (MP) and non-structural (NS) proteins. Whilst these were originally consistent with the different HA lineages circulating, multiple reassortment events have resulted in varying combinations of internal gene segment composition (Watson et al., 2015). This has not been investigated in England since 2014 and, given the increased diversity of SwIAVs observed in European pigs in recent years (Chastagner et al., 2020, Chepkwony et al., 2021, Encinas et al., 2022, Graaf-Rau et al., 2023), as well as sporadic human infections with SwIAVs (termed “variant” (v) influenza A virus) (Cogdale et al., 2024, Deng et al., 2020, Zhu et al., 2016, Freidl et al., 2014, Hennig et al., 2022, Rovida et al., 2017), there was a need for further characterisation of SwIAV genetic diversity within the pig populations in England. Consequently, in this study we describe the genetic diversification of H1 SwIAVs in England through whole-genome sequencing (WGS) analysis of clinical samples collected through passive surveillance between 2014 and 2021, and characterise the genetic diversity observed therein.

## Methods

### Sample screening and sample selection

Clinical samples for the study were obtained through the Government-funded GB swine influenza passive surveillance programme. Under this, samples (nasal swabs from live pigs or homogenates of trachea, tonsil and lung tissue obtained post-mortem) from pigs with respiratory signs and/or respiratory pathology are submitted for testing from the GB scanning surveillance network (Mollett et al., 2023, DEFRA, 2024b). Total RNA was extracted from these samples using the MagMAX™ CORE nucleic acid purification kit (ThermoFisher Scientific) according to the manufacturer’s instructions. The resulting RNA was then screened for SwIAV viral RNA (vRNA) to aid in sample selection using a reverse transcriptase PCR (RRT-qPCR targeting the viral M gene (Nagy et al., 2021, Slomka et al., 2010). Following a positive detection (Cq <=36), samples were analysed by whole genome sequencing (WGS).

### Whole Genome Sequencing

Extracted total RNA was utilised to amplify the viral RNA and generate double-stranded cDNA using sequence-independent single-primer amplification (SISPA) (Lewandowski et al., 2019) and purified using AMPure beads (Beckman Coulter, Brea, CA, USA). Sequencing library preparation was performed using the Nextera DNA Library Prep Kit (Illumina, Cambridge, MA, USA) and sequenced using the NextSeq System (Illumina). All kits were used as per the manufacturer’s instructions. Raw sequencing reads were assembled using a custom script as described previously (Byrne et al., 2023). All generated sequences were deposited in NCBI (Table S1).

### Phylogenetic analysis and clade classification

All publicly available European SwIAV sequences from 2006-2021 were obtained from GISAID EpiFlu, NCBI Virus and ESNIP3 (Watson et al., 2015) and combined with the data generated in this study to provide the necessary resolution of European and English viruses. Duplicate sequences were removed using seqkit v2.0.0 (Shen et al., 2016). Sequences were aligned using mafft v7.487 (Katoh and Standley, 2013) and manually trimmed to the open reading frame using AliView (Larsson, 2014). Clade classification of HA sequences was carried out via the bacterial and viral bioinformatics resource centre (BV-BRC) using the sub-species classification tool set to global swine H1 nomenclature (Anderson et al., 2016).

Phylogenetic trees for each gene segment were inferred using the maximum likelihood approach in IQ-Tree v2.1.4 (Minh et al., 2020) with ModelFinder to identify the best fit model of molecular evolution, 1,000 ultrafast bootstraps and subsampled using PARNAS (Markin et al., 2023b) to display at least 95% of diversity. Phylogenetic trees and associated matrices were visualized using R version 4.1.1 with the libraries Tidyverse (Wickham et al., 2019), ggtree (Xu et al., 2022), phytools (Revell, 2024) and treeio (Wang et al., 2020).

### Defining HA clades

New clades were defined by sharing of a common node and monophyly within swine, statistical support (greater than or equal to 90% SH-like approximate likelihood ratio test (Guindon et al., 2010) and greater than or equal to 75% ultrafast bootstrap (Hoang et al., 2018). The octoFLU pipeline was used to confirm classifications or identify areas that could not be classified with current approaches (Chang et al., 2019). As the octoFLU pipeline was generated to classify IAV in pigs in the United States, there is a revised reference dataset that can be used to classify whole genome sequence data for IAV in pigs in Europe at https://github.com/flu-crew/octoFLU. Within- and between-clade average pairwise distance (APD) was calculated using MEGACC v10.2.6 (Kumar et al., 2018) and threshold of within-clade mean nucleotide distance of below 5% and between clade-difference above 5% for HA and NA (Anderson et al., 2016) (Table S2).

### Defining genotypes

Genotypes were defined by identifying all English taxa and designating groups based on monophyly and paraphyly. The defined groups were labelled A-G, dependent on how many groups were present per gene, with A having the greatest number of representatives (Lycett et al., 2020). Genotypes were numbered 1-24 based on the letter combination for all genes (Table S3). Within- and between-clade APD was calculated for each group MEGACC v10.2.6 (Kumar et al., 2018) with the aim of having within clade mean distance of below 5% and between clade differences above 5% for HA and NA (Anderson et al., 2016). For the remaining gene segments, these thresholds were lowered to 3% (Table S2).

## Results

Between 2014-2021, 368 samples were screened using RT-PCR, with 170 samples from 82 independent submissions being positive for IAV. A total of 144 samples were selected for WGS, based on a Cq value ≤36 of which 82 (56%), spanning 71 submissions, yielded genomic data allowing for subtype determination based on HA and NA sequence data. From these samples, 78% (n=64/82) produced genomic data for all gene segments, formulating the dataset for genotyping. Distribution across the country varied but aligned with distribution of holdings and pig population density ((DEFRA, 2024a)). Positive submissions were derived from counties across England (Figure 1A, Table 1).

**Figure 1A:**
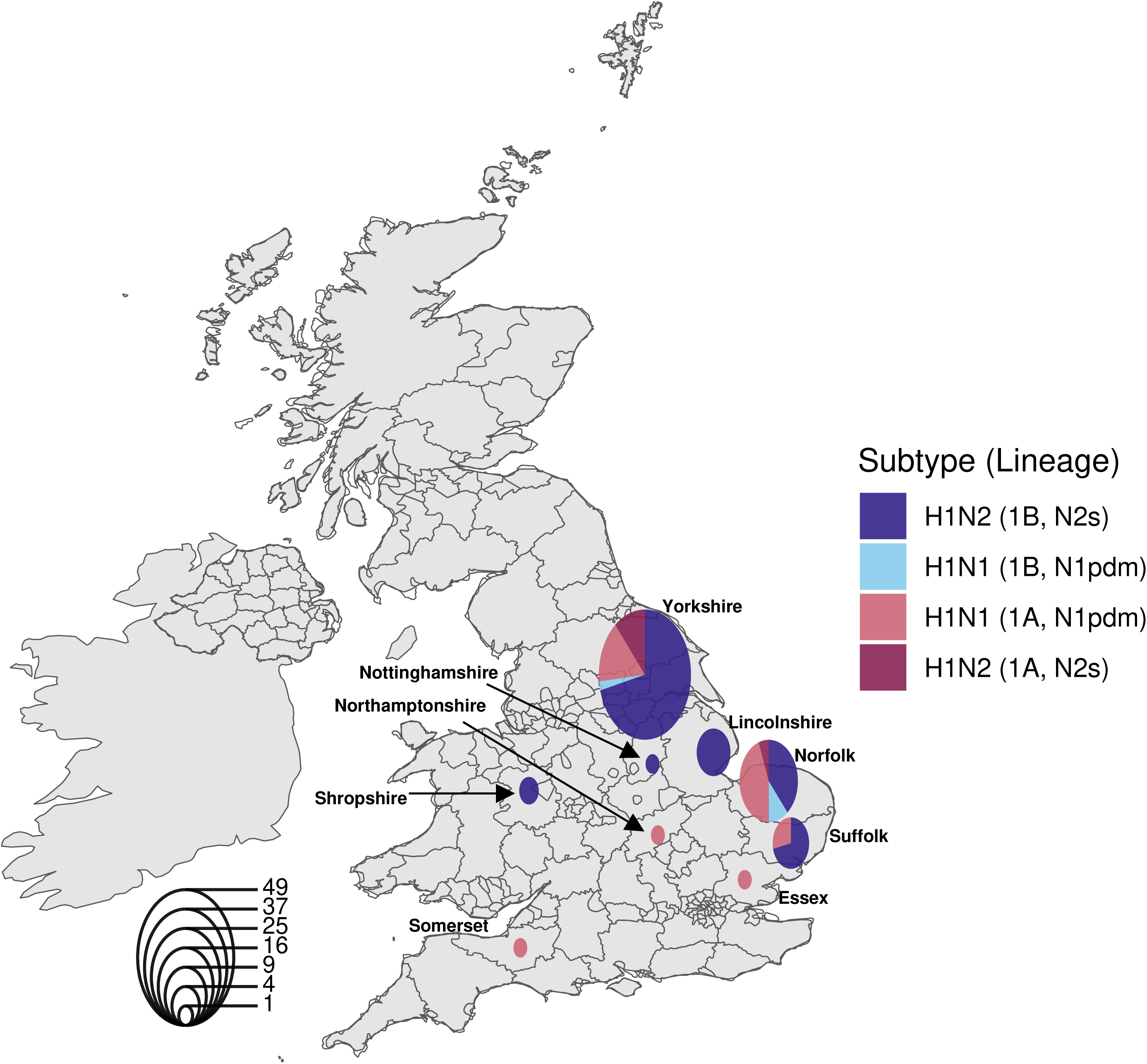
Geographical map of the United Kingdom depicting location of sample collection from 2014 to 2021 and identified with their haemagglutinin (HA) and neuraminidase (NA) nomenclature. The radius of the pie chart is indicative of the total number of positive samples observed per county and segments are coloured by subtype.

**Table 1:** Distribution of subtyped swine influenza samples and submissions per county.

| **County** | **Samples** | **Submissions** |
| --- | --- | --- |
| Yorkshire | 45 | 36 |
| Norfolk | 18 | 18 |
| Suffolk | 7 | 6 |
| Lincolnshire | 6 | 5 |
| Shropshire | 2 | 2 |
| Essex | 1 | 1 |
| Northamptonshire | 1 | 1 |
| Nottinghamshire | 1 | 1 |
| Somerset | 1 | 1 |
| Total | 82 | 71 |

### Subtyping

Each genome was evaluated using phylogenetic inference to determine the evolutionary lineage of the HA and NA glycoproteins. Classifications were confirmed using octoFLU (Chang et al., 2019). The HA of the 82 viruses subtyped belonged to two of the three global H1 lineages: 1A classical swine (32% [n=26]) and 1B human seasonal (68% [n=56]) (Figure 1A, Figure 2). Sequence data for the 1C viruses from the Eurasian avian lineage was not successfully sequenced, but this subtype was detected through the GB diagnostic and surveillance activities in 2014, 2015 and 2017 (S. Reid personal communication) and characterised using a molecular subtyping assay (Henritzi et al., 2016). H1 1A.3.3.2 and H1 1B.1.1.x clades were both detected in the pig dense regions of England (DEFRA, 2024a) with no indication of spatial segregation for either H1 clade within a particular region (Figure 1A). 1B.1.1.x clade viruses were the most prevalent across the duration of this study, accounting for the majority of subtyped viruses annually, with the exception of 2021 (Figure 1B). The 1A.3.3.2 clade viruses were comparatively less abundant, forming a minor proportion of the subtyped viruses throughout the timeframe of the study except for 2021 where 1A viruses were the most abundant. Prior to the 2014-2021 period, the 1B.1.1.x clade viruses were detected in continental Europe (France and Poland), but this HA group has, based on global data release, been exclusively restricted to GB since 2008. The English 1B lineage viruses fall within the 1B.1.1 clade with three distinct monophyletic groups (Figure 2). Conversely, the English 1A.3.3.2 viruses characterised in this study were found to be interspersed with European 1A.3.3.2 viruses (Figure 2) demonstrating either that the HA segment has evolved at the same rate between Europe and England or that there have been concurrent, likely bidirectional introductions of H1N1pdm09 human seasonal viruses between Europe and England.

**Figure 1B:**
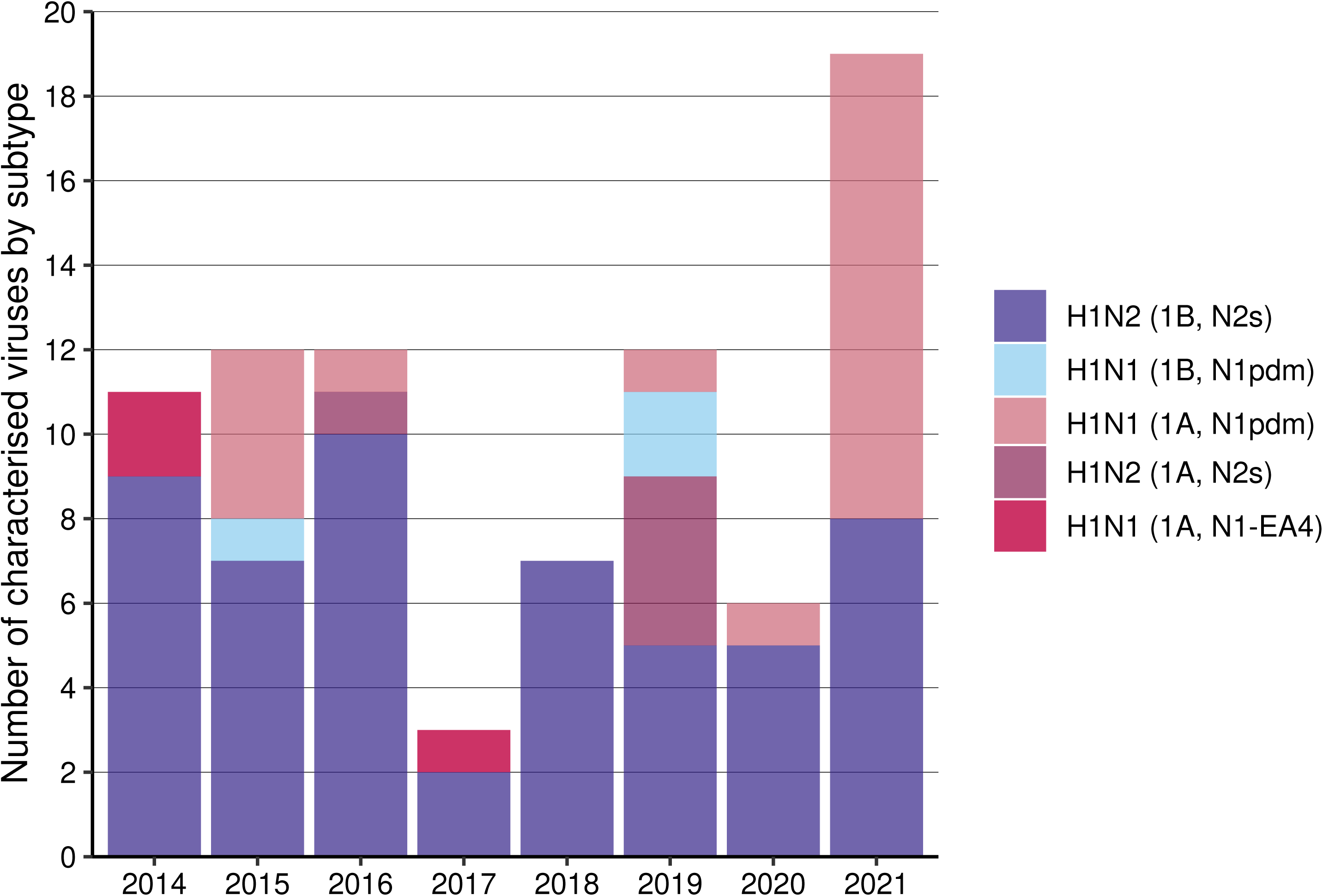
Number of SwIAV strains identified in England from 2014 to 2021, per year according to subtype.

**Figure 2:**
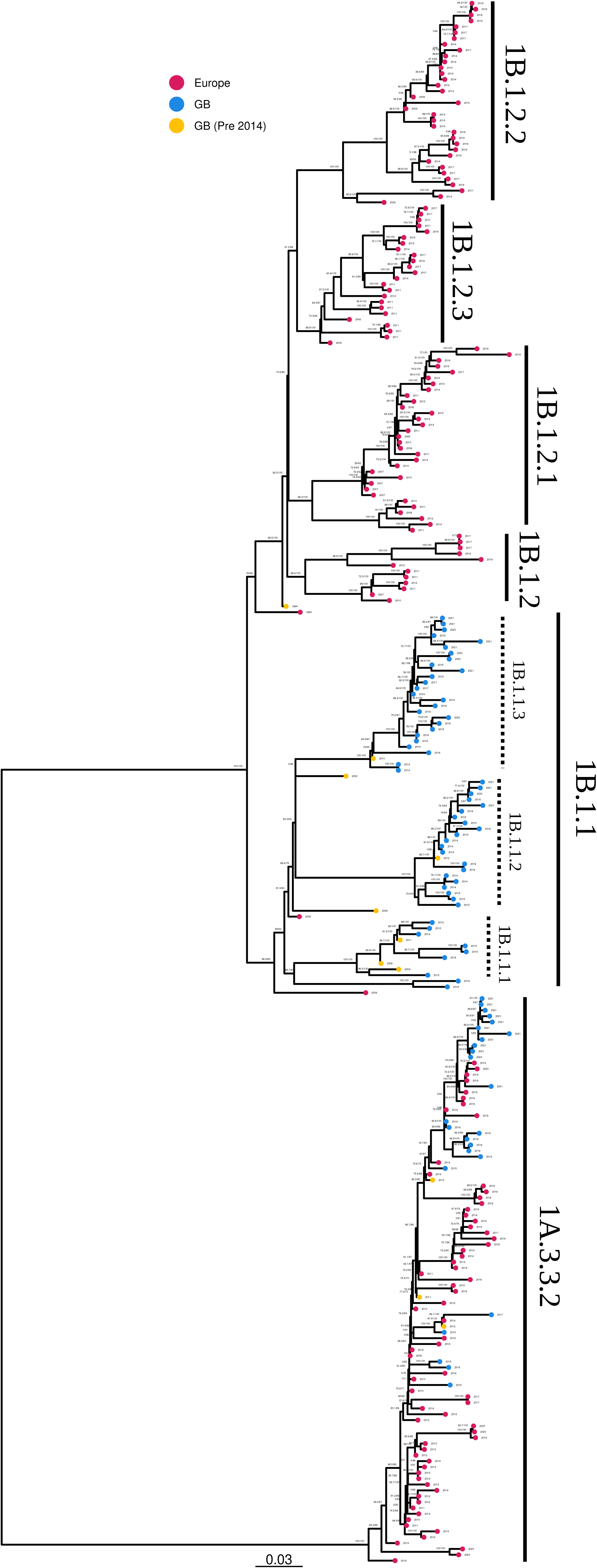
Maximum likelihood phylogeny of the haemagglutinin (HA) segment from European swine Influenza A viruses. The phylogeny is labelled by two of the major first order H1 lineages and 1B.1.1 clade is split by the proposed subclades detected in England. Tip points are coloured by sequences analysed in this study (Blue), publicly available GB sequences prior to 2014 (Yellow) and publicly available European sequences (Pink)

From the N1 genes characterised, 25% contained a H1N1pdm09 origin NA (n=21/82) (Figure 3). There were 3 occurrences of N1 segments detected within England during 2014 and 2017 which was classified as N1-Avian-4 (EA-4) (Figure 3) reflecting an avian-origin N1 that has persisted in pigs. The EA-4 has been previously detected in England in 1994, 2006 and 2013 (Figure 3). The N2 NA gene segments (n=58) were similar to the prototype virus, A/swine/Scotland/410440/1994 and therefore belong to a group considered Scotland-like (N2s) (Brown et al., 1994; Watson et al., 2015) (Figure 4). As with the 1B.1.1.x HA, the N2 segment characterised in English-origin viruses also had distinct monophyly, with no other European virus sequences being reported within that N2 clade (Figure 4). Conversely, the N1 pandemic segment clusters closely with other European strains and likely reflects the consistent introduction of 1A.3.3.2 viruses from human populations into swine with subsequent transmission in pigs in Europe and globally.

**Figure 3:**
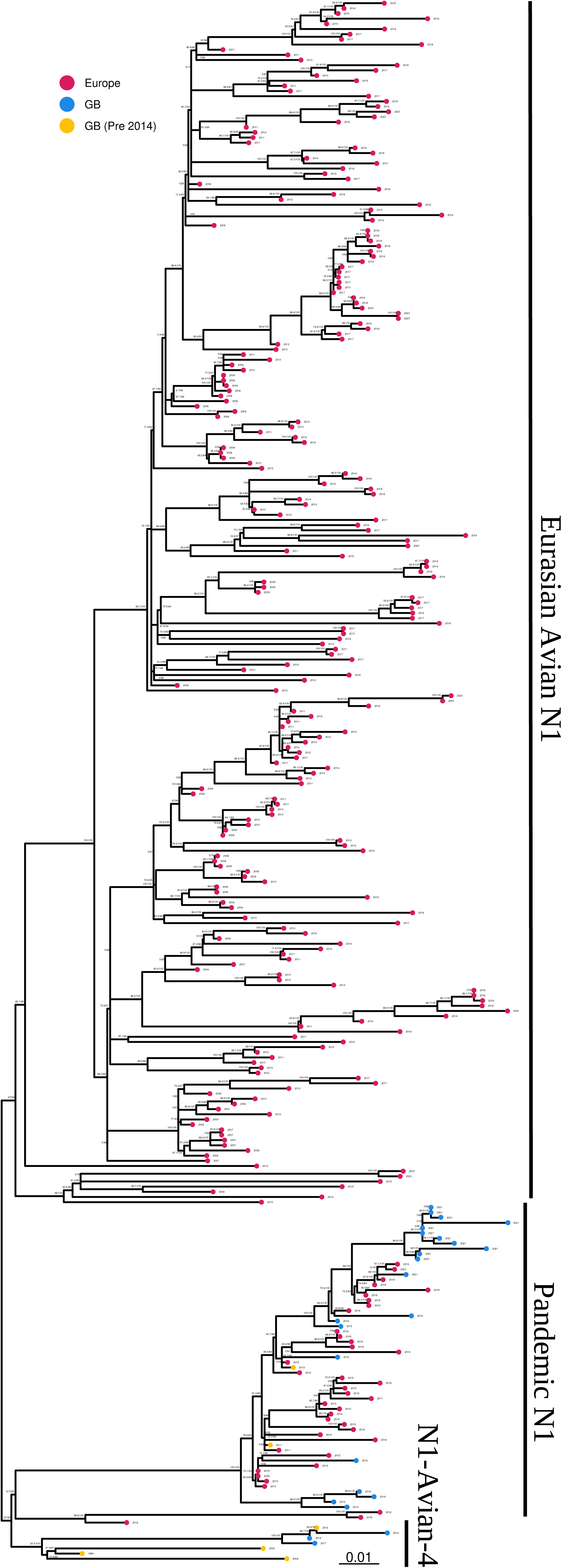
Maximum likelihood phylogeny of the N1 segment from European swine Influenza A viruses. The phylogeny is labelled by ancestral origin of the N1 segment. Tip points are coloured by sequences analysed in this study (Blue), publicly available GB sequences prior to 2014 (Yellow) and publicly available European sequences (Pink)

**Figure 4:**
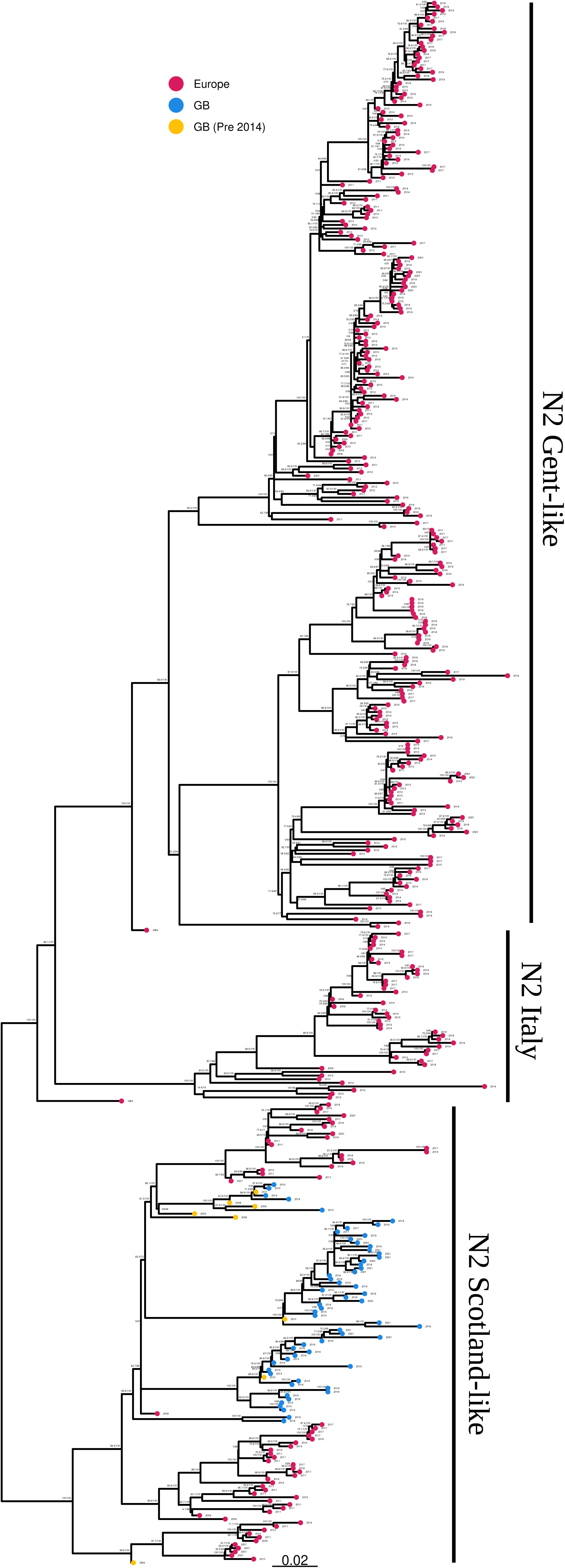
Maximum likelihood phylogeny of the N2 segment from European swine Influenza A viruses. The phylogeny is labelled by ancestral origin of the N2 segment. Tip points are coloured by sequences analysed in this study (Blue), publicly available GB sequences prior to 2014 (Yellow) and publicly available European sequences (Pink)

To quantify diversity in the glycoproteins within English SwIAVs, a set of thresholds were used to define clades with the aim of having within clade APD below 5% and between clade APD above 5%. APD within the English 1B.1.1 HA genes sequenced for this study was 9.8% (Table 3). Given this divergence, statistical support within the phylogeny, and consistent detections across multiple years within England, it is proposed that the 1B.1.1 clade divided into 3 sub-groups: 1B.1.1.1, 1B.1.1.2 and 1B.1.1.3 (Figure 2). The three groups were monophyletic, and following the division, the within- and between-group APD (Table S2) fell within the criteria established for naming SwIAV HA clades (Anderson et al. 2016; Anderson et al. 2021). The APD of the 1A.3.3.2 HA genes were 3.8% (Table 3) with no evidence to define different groups within these data. The newly defined clade classifications were applied to the 56 identified 1B.1.1 viruses, resulting in the following distribution: seven sequences in 1B.1.1.1, 21 in 1B.1.1.2, and 26 in 1B.1.1.3. Two additional viruses that were collected in 2018 and 2019 did not match the specific criteria of any group and remained under the broader 1B.1.1 category. APD within the N1 segments of H1N1pdm09 origin and EA-4 were 3.8% and 1.6%, respectively, providing no evidence to define different groups within these data. However, the N2s APD was found to be 8.3% indicating that new groups should be defined. As this NA lineage is widely distributed across Europe and not unique to England, new definitions were not proposed based on these data.

Molecular subtyping revealed that H1N2 and H1N1 subtypes predominated and cocirculated between 2014 and 2021; however, the evolutionary origin of the HA and NA segments varied. Most H1 HA subtypes were from the 1B.1.1.x clades and were paired with N2s (n=53/82 of sequenced samples). H1 HA 1A.3.3.2 clade viruses were paired with an N1 (H1N1pdm09 origin) (n=18/82 of sequenced samples). Reassortant viruses were sporadically detected and included the 1B.1.1.1 and 1B.1.1.2 HA paired with N1 (H1N1pdm09 origin) n=3/82 samples, and 1A.3.3.2 HA paired with N2s n=5/82 samples as well as three viruses incorporating 1A.3.3.2 HA paired with N1 (EA-4) (Figure 1A, Figure 1B).

The 1A.3.3.2 H1N1 and 1B.1.1.x H1N2 subtypes exhibited widespread distribution across multiple counties in England, indicating minimal spatial segregation (Figure 1A). In contrast, reassortant viruses, 1B.1.1.1 paired with N1 (H1N1pdm09 origin) and 1A.3.3.2 H1 paired with N2, were confined to high pig density regions in Yorkshire and Norfolk, while reassortant viruses containing 1B.1.1.2 paired with N1 (H1N1pdm09 origin) and 1A.3.3.2 with EA-4 NA segment detections restricted to samples from pigs in Yorkshire (Figure 1A). The 1B.1.1.x H1N2 was detected consistently across all study years, while pandemic-origin H1N1 was not observed in 2014, 2017, or 2018 (Figure 1B). The 1A.3.3.2 HA gene, associated with the EA-4 NA segment, was present in 2014 and 2017. 1B.1.1.1 coupled with N1 (H1N1pdm09 origin) and 1B.1.1.2 paired with N1 (H1N1pdm09 origin) were identified in 2019 and 2015 respectively. Finally, 1A.3.3.2 H1 paired with N2s appeared in 2016 and 2019 (Figure 1B).

### Genotyping

All viruses analysed contained genes of H1 1A.3.3.2 origin (Table 2, Supplementary Figure 1). The internal genes PB2, PB1, PA, NP and NS displayed >3% nucleotide differences, with MP having only 2.9% differences (Table 3). Based on pre-defined genetic divergence criteria, twenty-four genotypes were described from the 64 whole genome sequences and were given the genotype nomenclature 1-24 (Figure 5; Table S1). Of the 24 genotypes, 11 were only detected once across the study (3, 5, 10, 11, 13, 14, 15, 16, 19, 21 and 22)

**Figure 5:**
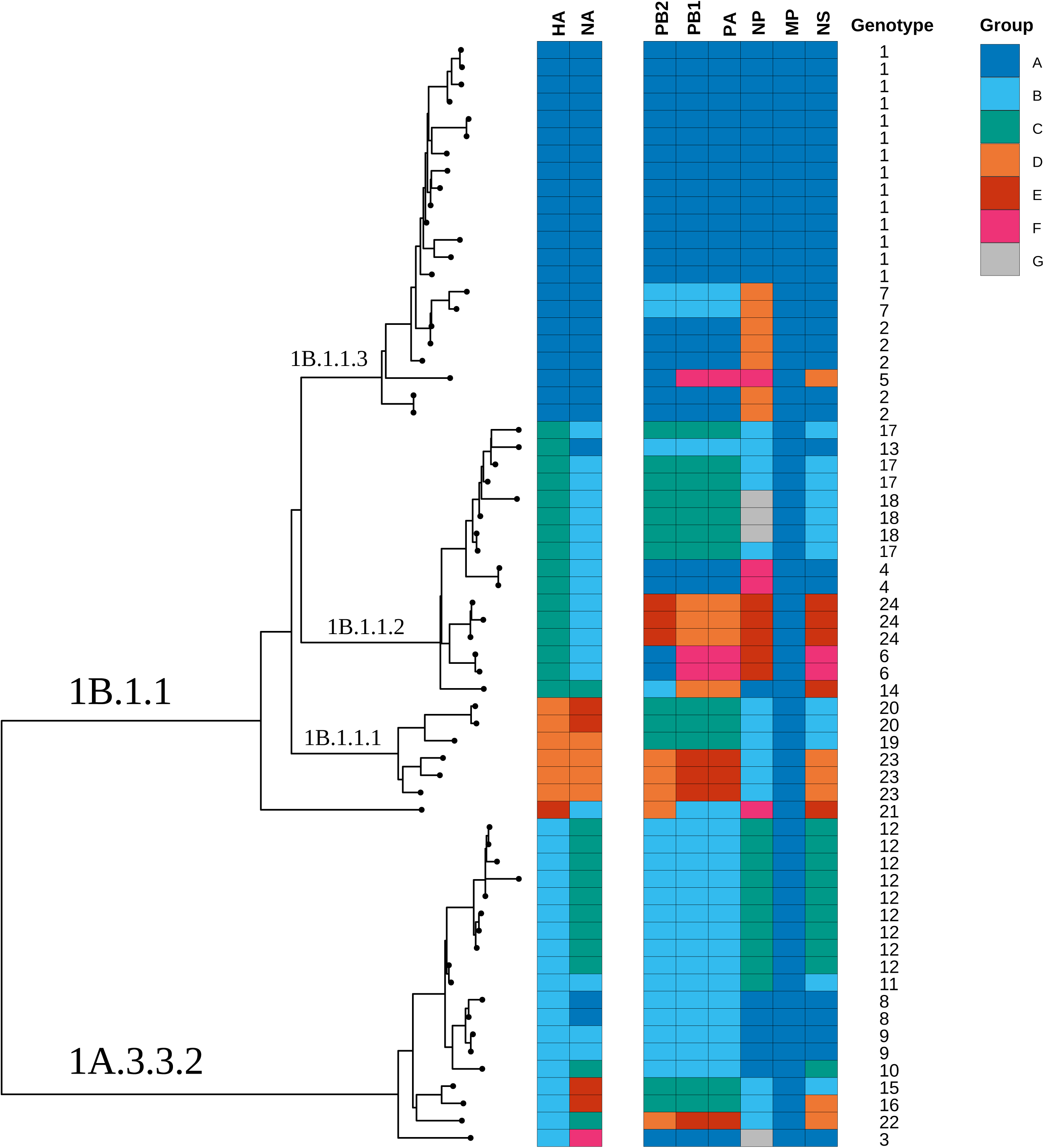
Maximum likelihood phylogeny of the England HA genes obtained in this study combined with a matrix displaying the internal gene constellations and associated genotype (1-24) nomenclature for each virus. The group represents the letter combinations used to define genotypes (Table S1). The matrix is coloured depending on which group the segment was allocated using average pairwise distance (Table S1, Table S2).

**Table 2:** Genomic constellation of all viruses genotyped using whole genome sequencing.

| Glycoproteins |  |  | Internal Genes |  |  |  |  |  | # | % |
| --- | --- | --- | --- | --- | --- | --- | --- | --- | --- | --- |
| Subtype | HA | NA | PB2 | PB1 | PA | NP | MP | NS |  |  |
| H1N2 | 1B.1.1 | N2s | pdm | pdm | pdm | pdm | pdm | pdm | 42 | 65.6 |
| H1N1 | 1B.1.1 | pdm | pdm | pdm | pdm | pdm | pdm | pdm | 03 | 4.7 |
| H1N1 | 1A.3.3.2 | pdm | pdm | pdm | pdm | pdm | pdm | pdm | 13 | 20.3 |
| H1N1 | 1A.3.3.2 | EA-4 | pdm | pdm | pdm | pdm | pdm | pdm | 01 | 1.6 |
| H1N2 | 1A.3.3.2 | N2s | pdm | pdm | pdm | pdm | pdm | pdm | 05 | 7.8 |

**Table 3:** Within group average pairwise distance (APD) of each gene segment for whole genome sequencing of English samples.

| Gene Segment | APD (%) |
| --- | --- |
| HA (1A.3.3.2) | 3.8 |
| HA (1B.1.1) | 9.8 |
| N1 | 3.8 |
| (H1N1pdm09) |  |
| N1 (EA-4) | 1.6 |
| N2 (N2s) | 8.3 |
| PB2 | 4.2 |
| PB1 | 3.8 |
| PA | 3.9 |
| NP | 3.6 |
| MP | 2.9 |
| NS | 4.0 |

Within the 1B.1.1.x clades, 14 genotypes were identified, with a further single genotype in the wider 1B.1.1 clade as it did not meet the criteria for subgroup designation. The 1B.1.1.3 clade includes genotypes 1, 2, and 7, with genotype 1 (n=14) being predominant, and differing from genotype 2 (n=5) only in NP (Figure 5). Genotype 7 (n=2) featured PB2, PB1, and PA segments from group B, typically associated with 1A.3.3.2 viruses (Figure 5).The 1B.1.1.2 clade includes genotypes 4, 6, 17, 18, and 24. Genotypes 4 (n=2) and 6 (n=2) exhibited gene differences in PB1, PA, NP, and NS (Figure 5). Genotypes 17 (n=4) and 18 (n=3), differing only in NP, were identified in multiple years, with genotype 17 observed in 2021 and genotype 18 in 2016 (Figure 6B, Table S1). Genotype 24 appeared three times in 2014 (Figure 6B, Table S1). The 1B.1.1.1 clade includes genotypes 20 (n=2) and 23 (n=3), which differ in NA, PB2, PB1, PA, and NS (Figure 5). Genotype 20 was detected in 2019, while genotype 23 was found in 2015 (Figure 6B, Table S1).

The 1A.3.3.2 clade comprises genotypes 8, 9, and 12. Genotypes 8 (n=2) and 9 (n=2) are reassortant viruses carrying N2s instead of the typical N1 (H1N1pdm09 origin), differing only in the divergence of their N2 segments (Figure 5). Genotype 8 was observed in 2019, and genotype 9 was detected in 2019 (Figure 6B, Table S1). Genotype 12 (n=9), the second most frequently observed genotype in this study, differs from genotypes 8 and 9 by its NP and NS segments (Figure 5). It was detected multiple times across different years (Figure 6B, Table S1). These data show that counties which were sampled more often display the greater diversity of genotypes (Figure 6A).

**Figure 6A:**
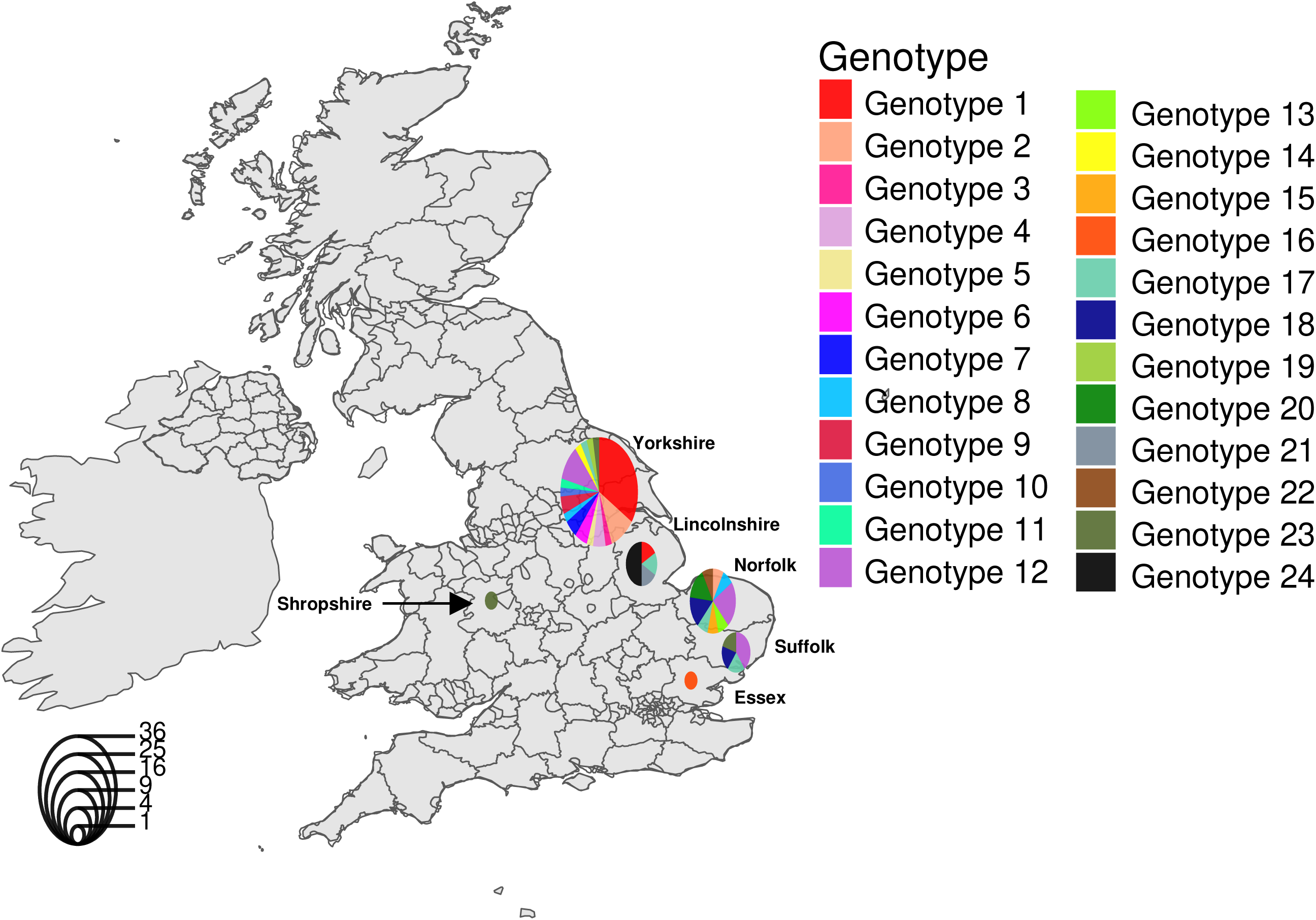
Geographical map of the United Kingdom depicting SwIAV genotypes detected in samples collected from 2014 to 2021. The radius of the pie chart is indicative of the total number of genotypes observed per county and segments are coloured by genotype.

**Figure 6B:**
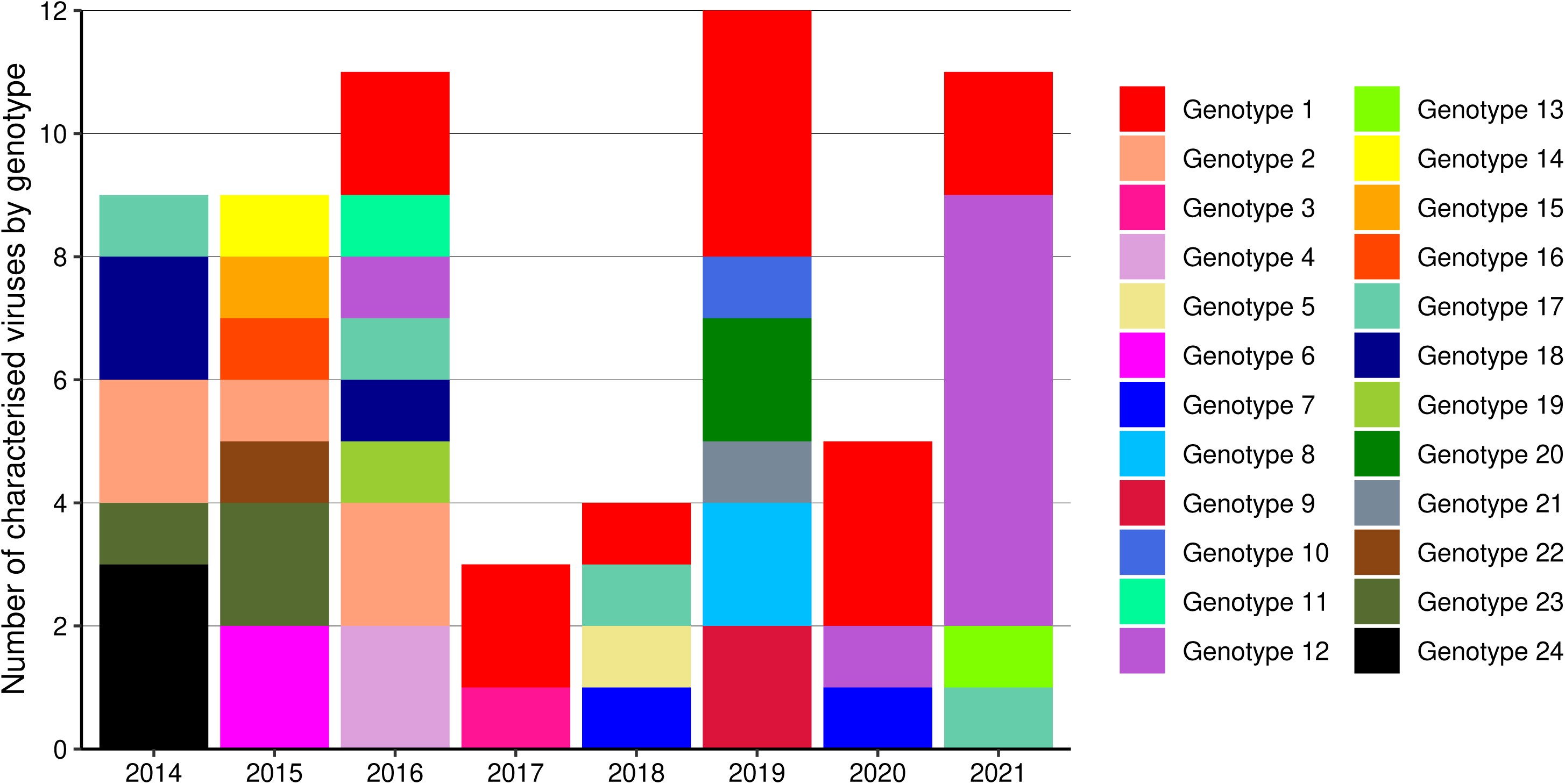
Number of SwIAV strains identified in England from 2014 to 2021, per year according to genotype. Genotypes occurring only once were omitted from the analysis.

## Discussion

A total of 82 swine SwIAVs were subtyped during this study, of which 64 were fully genotyped. This characterisation offers critical insights into the diversity of circulating SwIAV strains in England between 2014 and 2021 and facilitates an understanding of their genetic evolution over time. Subtyping revealed that the HA from 1B.1.1.x viruses paired with N2 neuraminidase and H1 1A.3.3.2 paired with N1 of H1N1pdm09 occurred most frequently indicating a preferential pairing and potential fitness peak whereby reassortants of alternative pairings are sporadic and seemingly unsuitable for sustained transmission in English pig populations. A similar trend has been observed in the U.S.A (Zeller et al., 2021, Hufnagel et al., 2023). The predominance of 1B.1.1.x contrasts with what has been reported across continental Europe where H1 1C lineage viruses predominated over a similar period (Henritzi et al., 2020). It may be that 1B.1.1.x viruses maintain a fitness advantage in English pigs, a finding that requires further investigation. None of the viruses characterised contained a HA derived from H1 1C lineage despite the ubiquity of this lineage in continental European pig populations (Brown, 2013, Li and Robertson, 2021). However, subtyping molecular assays did detect the presence of these viruses in 2014, 2015 and 2016. As more genomic data is produced it will be critical to assess 1C lineage viruses and whether they reassort with endemic 1A.3.3.2 and 1B.1.1.x viruses in England to determine impact in animal and public health.

A key outcome of this study was the further classification the 1B.1.1.x group of HA genes in England. This group has diverged into three monophyletic clades, exhibiting significant within-clade diversity (APD of 9.6%) and contemporaneous circulation from 2014 to 2021. Four distinct 1B.1.1 HA clades were identified; however, only three meet the criteria outlined by Anderson et al. (2016) for revising and designating new clades within the H1 global nomenclature system. It is proposed that subgroups 1B.1.1.1, 1B.1.1.2, and 1B.1.1.3 are adopted to better define this genetically diverse group. Clade 1B viruses not falling within these subclades retained 1B.1.1 designation. Similarly, mounting evidence suggests that the N2 lineage also meets the criteria for defining new genetic groups. All N2 viruses in this study formed discrete monophyletic clades, indicating independent evolution within English pig populations. The APD of 8.3% within the N2 group highlights the substantial genetic diversity present in this neuraminidase segment. This issue appears to be Europe-wide, necessitating a revision of the genetic classification of the neuraminidase gene, akin to efforts in North America (Zeller et al., 2021, Hufnagel et al., 2023). Such an approach would enhance our ability to monitor reassortment, identify emerging viruses, and improve communication between animal and public health sectors.

The use of complete genomic data for this study revealed that despite the variation in viral glycoproteins, all viruses detected contained a full complement of remaining genes derived from the H1 1A.3.3.2 viruses introduced to England in 2009. This gene constellation is often reported globally (Henritzi et al., 2020, Danilenko et al., 2021, Chiapponi et al., 2021, Richard, 2023, Kim et al., 2024, Nelson and Vincent, 2015) and it was not surprising that the viruses characterised had genetic components derived from H1N1pdm09 evolutionary lineage. Furthermore, there is evidence from multiple countries that spillover of human seasonal viruses into pig populations occurs sporadically, leading to the continued circulation of H1N1pdm09 associated genes (Junqueira et al., 2023, Nelson and Vincent, 2015, Kim et al., 2024). This is likely occurring in England but requires further investigation. Similar studies in Europe report Eurasian avian gene segments, originating from H1 1C viruses, mixing with H1 1A.3.3.2 genes to form multiple genotypes or indeed, create genotypes with wholly EA-origin internal genes (Chiapponi et al., 2021, Henritzi et al., 2020, Danilenko et al., 2021, Richard, 2023). Determining how reassortment to acquire H1N1pdm09 genes affects phenotype is a critical issue as these reassortment events may drive evolution of the envelope glycoproteins (Zeller et al., 2021). The acquisition of these gene segments may affect replicative fitness or transmission phenotypes (Thomas et al., 2024) and therefore impact zoonotic risk of endemically circulating SwIAVs in England.

The detection of greater genetic variability in SwIAV in England compared to previous studies necessitated genotypic classification aligned with established methods of genetic differentiation (Anderson et al., 2016, Byrne et al., 2023, Lycett et al., 2020). Using this approach, 24 genotypes were identified, contrasting with the five determined through traditional lineage-based genotyping focused on glycoproteins. Genotype 1 was the most prevalent among the 1B.1.1.x clade and was detected annually from 2016, suggesting gene combinations that confer a transmission advantage in pigs. For the 1A.3.3.2 clade, Genotype 12 was the most common, initially detected in 2016 and reappearing in 2020 and 2021. Its absence in 2017–2019 may reflect under-sampling or a later reintroduction into swine from humans. To fully understand these dynamics, additional influenza virus sequences from these periods are required. Moreover, the lack of a regular time series across premises hinders the identification of persistent genotypes and their broader distribution. Certain genetic constellations were retained across genotypes, warranting further investigation into the mechanisms influencing the maintenance and selection of specific genetic combinations. The implications for viral fitness and persistence in pig herds remain unclear.

In this study, the majority of samples were collected along the eastern side of England, where pig density is highest, particularly in East Anglia and the Northeast (DEFRA, 2024a). Minimal spatial segregation was observed between the two main subtypes, H1 1B.1.1.x paired with N2 and H1 1A.3.3.2 paired with N1 (H1N1pdm09). Most subtypes, including reassortants, were identified in Yorkshire and Norfolk, highlighting the need for enhanced surveillance in pig-dense regions to improve characterisation of endemic SwIAV. The EA4 N1 segment was primarily isolated to Yorkshire, potentially due to geographic restrictions or unsampled populations in other counties. Regions with higher representation in WGS data exhibited greater genotype diversity, underscoring the value of increased surveillance for improving SwIAV characterisation and risk assessments. Swine influenza is not a notifiable or reportable disease in England. This study was dependent on voluntary submissions to a government-funded passive surveillance programme, however veterinarians attending pigs can also use commercial laboratories for swine influenza diagnosis or may not test at all. Therefore, the Government surveillance scheme only detects a subset of active swine influenza infections and does not detect subclinical infections in pigs. The sample throughput and subsequent sequence data obtained was not equal in different years of this study so it is difficult to interpret changes in the number of any given genotype circulating in a particular year but will most likely be linked to disease burden. Vaccination plays a significant role in controlling influenza in pigs, but its effectiveness is limited by challenges in strain selection and infrequent updates to antigen composition (Everett et al., 2021). Monitoring virus evolution will help to increase vaccine efficacy and potentially reduce disease burden.

Swine-origin viruses containing genes from the 1A.3.3.2 clade have been sporadically detected in humans (Deng et al., 2020, Zhu et al., 2016, Freidl et al., 2014, Rovida et al., 2017, Hennig et al., 2022). Recently, the first reported case of influenza A(H1N2)v from the 1B.1.1.1 clade was identified in the United Kingdom (Cogdale et al., 2024). Notably, representative viruses with similar genetic compositions exhibited poor cross-reactivity in haemagglutination inhibition assays with human seasonal vaccine viruses, suggesting that current vaccines are unlikely to offer cross-protection. Fortunately, no onward transmission to contacts was observed in this case (Cogdale et al., 2024). Although not genetically characterised in this study, H1 1C lineage viruses are sporadically detected in England and have been associated with variant cases in Europe. Should these SwIAVs acquire the ability to replicate efficiently and transmit between humans, they would pose a significant pandemic risk. Antigenic assessment of such strains is essential to evaluate cross-reactivity with viruses contributing to human population immunity.

In conclusion, alongside the need for enhanced surveillance strategies to improve the resolution of SwIAVs in England, it is critical that WGS is deployed wherever possible to detect reassortment patterns that may influence the transmission phenotype of viruses, even if the HA and NA genes are similar to prior detections. To that end, including rapid and cost-effective technologies, such as nanopore technology, may help to provide a breadth of relevant genomic information and may also be feasible to carry out as a frontline diagnostic procedure (Rambo-Martin et al., 2020, Vereecke et al., 2023).

## Acknowledgments

The authors would like to acknowledge the originating and submitting laboratories of the sequences from GISAID’s EpiFlu Database upon which this research is based, and analyses described in the text. All submitters of the data may be contacted directly via the GISAID website (https://www.gisaid.org). The analyses described in this work were conducted using the Scientific Computing Environment at the Animal and Plant Health Agency.

The authors would also like to acknowledge and thank the pig farmers and veterinarians across the UK without whom the sample collection and subsequent analysis would not be possible.

---

**Supplementary Table 1:** Strain names and associated GISAID accession number

**Supplementary Table 2:** within- and between clade average pairwise distance for each gene of the English viruses characterised by WGS.

**Supplementary Table 3:** Genotyping assignment table depicting groups assigned, the overall combination of groups and final genotype

**Supplementary Figure 1:** Maximum likelihood Phylogeny of gene segments from European swine Influenza A viruses and the 64 English samples sequenced in this study. The phylogeny is labelled by gene. Tip points are coloured by the region or time when defining UK samples collected prior to this current studies dataset.

