## Supplementary figures and images for "Diverse Genomic Landscape of Swine Influenza A Virus in England (2014–2021)"

### Figure S1 - Internal Genes.pdf

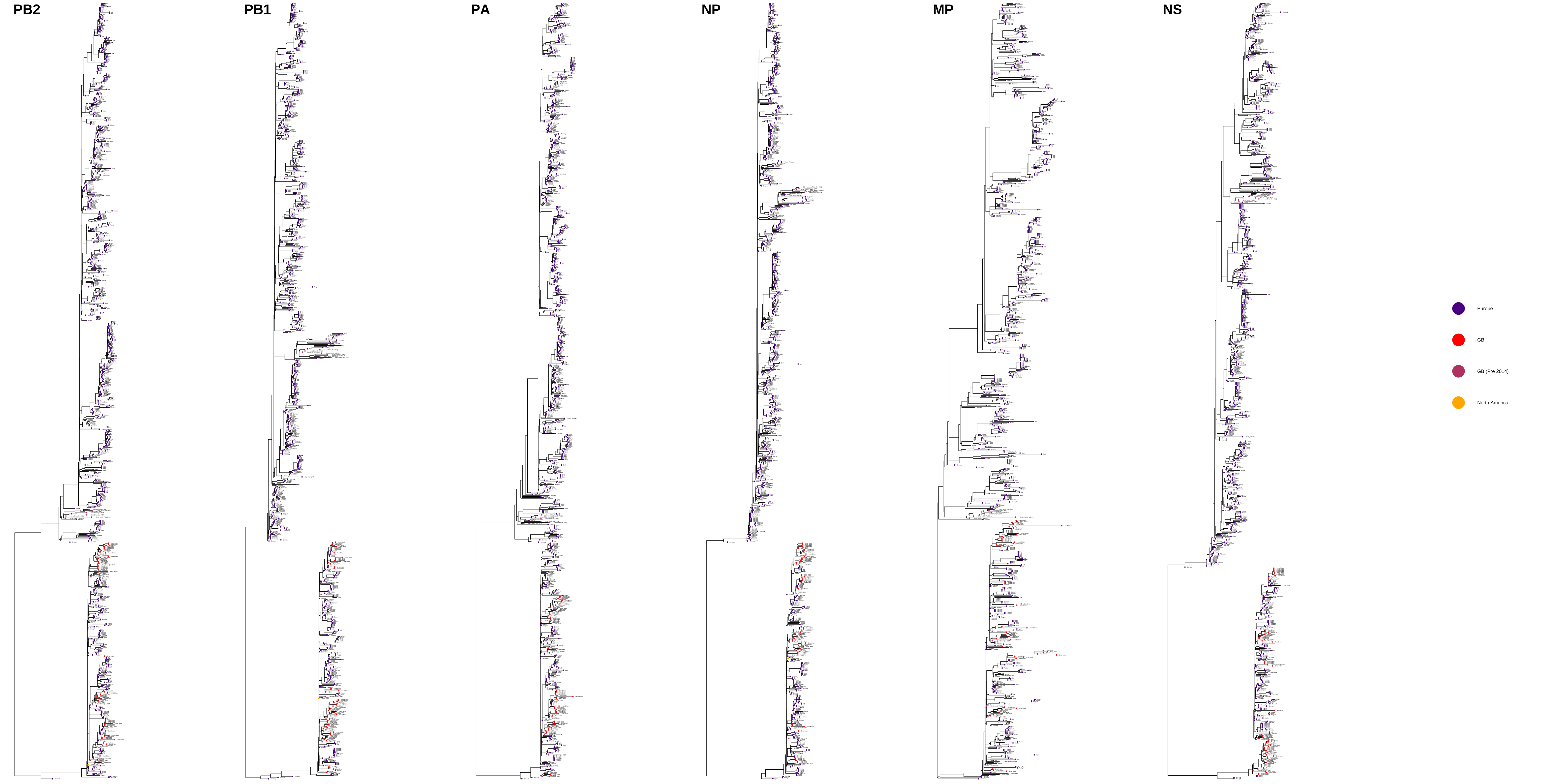
