## Supplemental Table 1 for "Diverse Genomic Landscape of Swine Influenza A Virus in England (2014–2021)": Table S1 - Isolate ID.pdf

| strain | Accession Number |
|---|---|
| A/swine/England/102081/2021 | EPI_ISL_18042673 |
| A/swine/England/102076/2021 | EPI_ISL_18042672 |
| A/swine/England/075667/2019 | EPI_ISL_18042657 |
| A/swine/England/092944/2020 | EPI_ISL_18042667 |
| A/swine/England/023799/2020 | EPI_ISL_18042637 |
| A/swine/England/023795/2020 | EPI_ISL_18042636 |
| A/swine/England/062942/2018 | EPI_ISL_18420064 |
| A/swine/England/180277/2017 | EPI_ISL_18042689 |
| A/swine/England/053048/2017 | EPI_ISL_18042652 |
| A/swine/England/072591/2019 | EPI_ISL_18421717 |
| A/swine/England/042792/2016 | EPI_ISL_18042646 |
| A/swine/England/044988/2016 | EPI_ISL_18378252 |
| A/swine/England/03180/2019 | EPI_ISL_18042644 |
| A/swine/England/079389/2019 | EPI_ISL_18421854 |
| A/swine/England/151782/2016 | EPI_ISL_18042682 |
| A/swine/England/151720/2016 | EPI_ISL_18042681 |
| A/swine/England/132457/2015 | EPI_ISL_18042675 |
| A/swine/England/091738/2014 | EPI_ISL_18424258 |
| A/swine/England/091736/2014 | EPI_ISL_18042664 |
| A/swine/England/051216/2017 | EPI_ISL_18378690 |
| A/swine/England/153554/2016 | EPI_ISL_18042684 |
| A/swine/England/153553/2016 | EPI_ISL_18042683 |
| A/swine/England/066871/2018 | EPI_ISL_18420262 |
| A/swine/England/143740/2015 | EPI_ISL_18042680 |
| A/swine/England/143739/2015 | EPI_ISL_18042679 |
| A/swine/England/064519/2018 | EPI_ISL_18420074 |
| A/swine/England/263079/2020 | EPI_ISL_18042701 |
| A/swine/England/078337/2019 | EPI_ISL_18042658 |
| A/swine/England/226704/2019 | EPI_ISL_18042693 |
| A/swine/England/016993/2019 | EPI_ISL_18042634 |
| A/swine/England/016994/2019 | EPI_ISL_18367865 |
| A/swine/England/082444/2019 | EPI_ISL_18422466 |
| A/swine/England/175093/2016 | EPI_ISL_18424267 |
| A/swine/England/237719/2021 | EPI_ISL_18424269 |
| A/swine/England/237596/2021 | EPI_ISL_18350490 |
| A/swine/England/237500/2021 | EPI_ISL_18350489 |
| A/swine/England/237498/2021 | EPI_ISL_18350488 |
| A/swine/England/238877/2021 | EPI_ISL_18424270 |
| A/swine/England/031764/2021 | EPI_ISL_18378245 |
| A/swine/England/031766/2021 | EPI_ISL_18350364 |
| A/swine/England/096411/2020 | EPI_ISL_18350473 |
| A/swine/England/043261/2016 | EPI_ISL_18350371 |
| A/swine/England/238434/2021 | EPI_ISL_18042698 |
| A/swine/England/040574/2015 | EPI_ISL_18378211 |
| A/swine/England/135933/2015 | EPI_ISL_18350478 |
| A/swine/England/142339/2015 | EPI_ISL_18350483 |
| A/swine/England/155224/2016 | EPI_ISL_18042685 |

| | |
|---|---|
| A/swine/England/204998/2018 | EPI_ISL_18370673 |
| A/swine/England/100227/2021 | EPI_ISL_18042671 |
| A/swine/England/096724/2014 | EPI_ISL_18042669 |
| A/swine/England/161192/2016 | EPI_ISL_18424260 |
| A/swine/England/091215/2014 | EPI_ISL_18042663 |
| A/swine/England/087377/2014 | EPI_ISL_18042662 |
| A/swine/England/045983/2016 | EPI_ISL_18042649 |
| A/swine/England/015824/2019 | EPI_ISL_18367709 |
| A/swine/England/219508/2019 | EPI_ISL_18424268 |
| A/swine/England/083412/2019 | EPI_ISL_18042661 |
| A/swine/England/140499/2015 | EPI_ISL_18350482 |
| A/swine/England/220198/2015 | EPI_ISL_18042692 |
| A/swine/England/130543/2015 | EPI_ISL_18042674 |
| A/swine/England/097085/2014 | EPI_ISL_18042670 |
| A/swine/England/028258/2014 | EPI_ISL_18370596 |
| A/swine/England/028256/2014 | EPI_ISL_18042640 |
| A/swine/England/026605/2014 | EPI_ISL_18042639 |
| A/Swine/England/239257/2021 | EPI_ISL_19675139 |
| A/Swine/England/238879/2021 | EPI_ISL_19675138 |
| A/Swine/England/238809/2021 | EPI_ISL_19675137 |
| A/Swine/England/228996/2020 | EPI_ISL_19675136 |
| A/Swine/England/161264/2016 | EPI_ISL_19675135 |
| A/Swine/England/143370/2015 | EPI_ISL_19675134 |
| A/Swine/England/133494/2015 | EPI_ISL_19675133 |
| A/Swine/England/125748/2015 | EPI_ISL_19675132 |
| A/Swine/England/104927/2021 | EPI_ISL_19675131 |
| A/Swine/England/103125/2021 | EPI_ISL_19675130 |
| A/Swine/England/103120/2021 | EPI_ISL_19675129 |
| A/Swine/England/090354/2014 | EPI_ISL_19675128 |
| A/Swine/England/087475/2014 | EPI_ISL_19675127 |
| A/Swine/England/081761/2018 | EPI_ISL_19675126 |
| A/Swine/England/062058/2018 | EPI_ISL_19675125 |
| A/Swine/England/033315/2021 | EPI_ISL_19675124 |
| A/Swine/England/033312/2021 | EPI_ISL_19675123 |
| A/Swine/England/004146/2018 | EPI_ISL_19675122 |
