## Supplementary Table 2 for "Diverse Genomic Landscape of Swine Influenza A Virus in England (2014–2021)": Table S2 - APD.pdf

| Gene | Group | APD (%) | | | | |
|---|---|---|---|---|---|---|
| | | Within Clade | Between Clade | | | |
| | | A | B | C | D | E |
| | A | 2.9 | | | | |
| PB2 | B | 2.4 | 5.3 | | | |
| | C | 1.7 | 4.4 | 4.6 | | |
| | D | 2.8 | 4.7 | 4.0 | 4.0 | |
| | E | 0.8 | 4.4 | 4.2 | 3.8 | 3.7 |

| Gene | Group | APD (%) | | | | | |
|---|---|---|---|---|---|---|---|
| | | Within Clade | Between Clade | | | | |
| | | A | B | C | D | E | F |
| | A | 2.7 | | | | | |
| | B | 2.2 | 3.2 | | | | |
| PB1 | C | 1.9 | 3.6 | 4.2 | | | |
| | D | 2.4 | 3.5 | 4.3 | 4.4 | | |
| | E | 1.5 | 3.8 | 4.4 | 4.7 | 4.9 | |
| | F | 2.2 | 2.6 | 3.5 | 3.6 | 3.6 | 4.0 |

| Gene | Group | APD (%) | | | | | |
|---|---|---|---|---|---|---|---|
| | | Within Clade | Between Clade | | | | |
| | | A | B | C | D | E | F |
| | A | 2.7 | | | | | |
| | B | 2.3 | 3.2 | | | | |
| PA | C | 1.9 | 3.3 | 3.4 | | | |
| | D | 2.1 | 4.0 | 4.1 | 4.4 | | |
| | E | 1.5 | 4.4 | 4.4 | 4.6 | 5.0 | |
| | F | 2.1 | 3.2 | 3.3 | 3.5 | 4.3 | 3.9 |

| Gene | Group | APD (%) | | | | |
|---|---|---|---|---|---|---|
| | | Within Clade | Between Clade | | | |
| | | A | B | C | D | E |
| | A () | 3.3 | | | | |
| HA | B | 3.8 | 24.4 | | | |
| | C | 3.5 | 13.1 | 25.8 | | |
| | D | 4.3 | 12.6 | 24.8 | 13.9 | |
| | E | N/A | 13.8 | 24.6 | 14.9 | 14.2 |

| Gene | Group | APD (%) | | | | | | |
|---|---|---|---|---|---|---|---|---|
| | | Within Clade | Between Clade | | | | | |
| | | A | B | C | D | E | F | |
| | A | 2.0 | | | | | | |
| | B | 2.3 | 3.6 | | | | | |
| NP | C | 1.0 | 3.9 | 4.1 | | | | |
| | D | 2.5 | 3.4 | 3.6 | 4.1 | | | |
| | E | 1.5 | 3.6 | 3.7 | 3.7 | 3.6 | | |
| | F | 2.9 | 4.2 | 4.2 | 4.4 | 4.1 | 4.3 | |
| | G | 2.7 | 3.9 | 4.1 | 4.4 | 3.9 | 4.4 | 3.4 |

| Gene | Group | APD (%) |
|---|---|---|

| | | | | | | | | |
|---|---|---|---|---|---|---|---|---|
| | | Within Clade | Between Clade | | | | | |
| | | | A | B | C | D | E | F |
| NA | A | 3.7 | | | | | | |
| | B | 4.4 | 11.7 | | | | | |
| | C | 2.8 | 64.2 | 64.2 | | | | |
| | D | 1.6 | 11.0 | 9.7 | 63.6 | | | |
| | E | 1.6 | 64.1 | 64.7 | 5.8 | 63.7 | | |
| | F | N/A | 64.7 | 64.4 | 11.0 | 64.6 | 10.6 | |

| Gene | Group | APD (%) | |
|---|---|---|---|
| | | Within Clade | Between Clade |
| MP | | | A |
| | A | 2.9 | |

| Gene | Group | APD (%) | | | | | | |
|---|---|---|---|---|---|---|---|---|
| | | Within Clade | Between Clade | | | | | |
| | | | A | B | C | D | E | F |
| NS | A | 2.2 | | | | | | |
| | B | 1.9 | 4.5 | | | | | |
| | C | 1.1 | 5.5 | 3.6 | | | | |
| | D | 2.6 | 5.5 | 3.8 | 3.9 | | | |
| | E | 2.9 | 5.7 | 4.1 | 4.0 | 4.4 | | |
| | F | 0.5 | 6.0 | 4.4 | 4.1 | 4.9 | 4.6 | |

G
