## Supplementary Table 3 for "Diverse Genomic Landscape of Swine Influenza A Virus in England (2014–2021)": Table S3 - Genotyping.pdf

| strain | PB2 | PB1 | PA | HA | NP | NA | MP | NS | Letter Combination | Genotype |
|---|---|---|---|---|---|---|---|---|---|---|
| A/swine/England/102081/2021 | A | A | A | A | A | A | A | A | AAAAAAAA | 1 |
| A/swine/England/102076/2021 | A | A | A | A | A | A | A | A | AAAAAAAA | 1 |
| A/swine/England/075667/2019 | A | A | A | A | A | A | A | A | AAAAAAAA | 1 |
| A/swine/England/092944/2020 | A | A | A | A | A | A | A | A | AAAAAAAA | 1 |
| A/swine/England/023799/2020 | A | A | A | A | A | A | A | A | AAAAAAAA | 1 |
| A/swine/England/023795/2020 | A | A | A | A | A | A | A | A | AAAAAAAA | 1 |
| A/swine/England/062942/2018 | A | A | A | A | A | A | A | A | AAAAAAAA | 1 |
| A/swine/England/180277/2017 | A | A | A | A | A | A | A | A | AAAAAAAA | 1 |
| A/swine/England/053048/2017 | A | A | A | A | A | A | A | A | AAAAAAAA | 1 |
| A/swine/England/072591/2019 | A | A | A | A | A | A | A | A | AAAAAAAA | 1 |
| A/swine/England/042792/2016 | A | A | A | A | A | A | A | A | AAAAAAAA | 1 |
| A/swine/England/044988/2016 | A | A | A | A | A | A | A | A | AAAAAAAA | 1 |
| A/swine/England/03180/2019 | A | A | A | A | A | A | A | A | AAAAAAAA | 1 |
| A/swine/England/079389/2019 | A | A | A | A | A | A | A | A | AAAAAAAA | 1 |
| A/swine/England/151782/2016 | A | A | A | A | D | A | A | A | AAAADAAA | 2 |
| A/swine/England/151720/2016 | A | A | A | A | D | A | A | A | AAAADAAA | 2 |
| A/swine/England/132457/2015 | A | A | A | A | D | A | A | A | AAAADAAA | 2 |
| A/swine/England/091738/2014 | A | A | A | A | D | A | A | A | AAAADAAA | 2 |
| A/swine/England/091736/2014 | A | A | A | A | D | A | A | A | AAAADAAA | 2 |
| A/swine/England/051216/2017 | A | A | A | B | G | F | A | A | AAABGFAA | 3 |
| A/swine/England/153554/2016 | A | A | A | C | F | B | A | A | AAACFBAA | 4 |
| A/swine/England/153553/2016 | A | A | A | C | F | B | A | A | AAACFBAA | 4 |
| A/swine/England/066871/2018 | A | F | F | A | F | A | A | D | AFFAFAAD | 5 |
| A/swine/England/143740/2015 | A | F | F | C | E | B | A | F | AFFCEBAF | 6 |
| A/swine/England/143739/2015 | A | F | F | C | E | B | A | F | AFFCEBAF | 6 |
| A/swine/England/064519/2018 | B | B | B | A | D | A | A | A | BBBADAAA | 7 |
| A/swine/England/263079/2020 | B | B | B | A | D | A | A | A | BBBADAAA | 7 |
| A/swine/England/078337/2019 | B | B | B | B | A | A | A | A | BBBBAAAA | 8 |
| A/swine/England/226704/2019 | B | B | B | B | A | A | A | A | BBBBAAAA | 8 |
| A/swine/England/016993/2019 | B | B | B | B | A | B | A | A | BBBBABAA | 9 |
| A/swine/England/016994/2019 | B | B | B | B | A | B | A | A | BBBBABAA | 9 |
| A/swine/England/082444/2019 | B | B | B | B | A | C | A | C | BBBBACAC | 10 |
| A/swine/England/175093/2016 | B | B | B | B | C | B | A | B | BBBBCBAB | 11 |
| A/swine/England/237719/2021 | B | B | B | B | C | C | A | C | BBBBCCAC | 12 |
| A/swine/England/237596/2021 | B | B | B | B | C | C | A | C | BBBBCCAC | 12 |
| A/swine/England/237500/2021 | B | B | B | B | C | C | A | C | BBBBCCAC | 12 |
| A/swine/England/237498/2021 | B | B | B | B | C | C | A | C | BBBBCCAC | 12 |
| A/swine/England/238877/2021 | B | B | B | B | C | C | A | C | BBBBCCAC | 12 |
| A/swine/England/031764/2021 | B | B | B | B | C | C | A | C | BBBBCCAC | 12 |
| A/swine/England/031766/2021 | B | B | B | B | C | C | A | C | BBBBCCAC | 12 |
| A/swine/England/096411/2020 | B | B | B | B | C | C | A | C | BBBBCCAC | 12 |
| A/swine/England/043261/2016 | B | B | B | B | C | C | A | C | BBBBCCAC | 12 |
| A/swine/England/238434/2021 | B | B | B | C | B | A | A | A | BBBCBAAA | 13 |
| A/swine/England/040574/2015 | B | D | D | C | A | C | A | E | BDDCACAE | 14 |
| A/swine/England/135933/2015 | C | C | C | B | B | E | A | B | CCCBBEAB | 15 |
| A/swine/England/142339/2015 | C | C | C | B | B | E | A | D | CCCBBEAD | 16 |
| A/swine/England/155224/2016 | C | C | C | C | B | B | A | B | CCCCBBAB | 17 |
| A/swine/England/204998/2018 | C | C | C | C | B | B | A | B | CCCCBBAB | 17 |
| A/swine/England/100227/2021 | C | C | C | C | B | B | A | B | CCCCBBAB | 17 |

| | | | | | | | | | | |
|---|---|---|---|---|---|---|---|---|---|---|
| A/swine/England/096724/2014 | C | C | C | C | B | B | A | B | CCCCBBAB | 17 |
| A/swine/England/161192/2016 | C | C | C | C | G | B | A | B | CCCCGBAB | 18 |
| A/swine/England/091215/2014 | C | C | C | C | G | B | A | B | CCCCGBAB | 18 |
| A/swine/England/087377/2014 | C | C | C | C | G | B | A | B | CCCCGBAB | 18 |
| A/swine/England/045983/2016 | C | C | C | D | B | D | A | B | CCCDBDAB | 19 |
| A/swine/England/015824/2019 | C | C | C | D | B | E | A | B | CCCDBEAB | 20 |
| A/swine/England/219508/2019 | C | C | C | D | B | E | A | B | CCCDBEAB | 20 |
| A/swine/England/083412/2019 | D | B | B | E | F | B | A | E | DBBEFBAE | 21 |
| A/swine/England/140499/2015 | D | E | E | B | B | C | A | D | DEEBBCAD | 22 |
| A/swine/England/220198/2015 | D | E | E | D | B | D | A | D | DEEDBDAD | 23 |
| A/swine/England/130543/2015 | D | E | E | D | B | D | A | D | DEEDBDAD | 23 |
| A/swine/England/097085/2014 | D | E | E | D | B | D | A | D | DEEDBDAD | 23 |
| A/swine/England/028258/2014 | E | D | D | C | E | B | A | E | EDDCEBAE | 24 |
| A/swine/England/028256/2014 | E | D | D | C | E | B | A | E | EDDCEBAE | 24 |
| A/swine/England/026605/2014 | E | D | D | C | E | B | A | E | EDDCEBAE | 24 |
